# Differential chromatin accessibility between pre- and post-natal stages highlights putative causal regulatory variants in pig skeletal muscle

**DOI:** 10.1101/2025.11.27.690935

**Authors:** Ervin Shishmani, Andrea Rau, Sarah Djebali, Emily L. Clark, Jordi Estellé, Valentino Palombo, Mariasilvia D’Andrea, Elisabetta Giuffra

## Abstract

Deciphering how chromatin dynamics shape the effects of regulatory variants on complex traits across development remains largely unexplored in farmed animals.

Focusing on porcine skeletal muscle, we integrated chromatin accessibility (ATAC-seq), molecular QTL (molQTL), and GWAS data to identify putative stage-specific regulatory variants influencing agronomically important traits.

We aggregated and analyzed 202 ATAC-seq libraries spanning prenatal (fetal) and postnatal (piglet) stages. We intersected consensus and differential peaks with cis-molQTLs from the Porcine Genotype-by-Tissue-by-Expression Atlas (PigGTEx) followed by functional enrichment analysis, promoter-focused functional annotation, integration with gene–trait pairs, and intersection with eGenes (i.e., genes whose expression is regulated by at least one cis-eQTL) to link variants previously associated with QTLs for production traits.

We identified 132,275 differentially accessible peaks (DAPs) distinguishing fetal and piglet muscle. Fetal DAPs were enriched in promoter and intergenic regions, whereas piglet DAPs were enriched in intronic regions, indicating a shift from transcriptional priming to postnatal regulatory refinement. Overlaps between accessible regions and molQTLs were highly significant (p < 0.001), with the strongest eQTLs predominating in the fetal stage (14 fetal vs. 4 piglet). Gene Ontology analysis of promoter-accessible eGenes revealed enrichment for RNA metabolism and chromatin organization in fetuses, and for muscle contraction and lipid metabolism in piglets. The intersection with colocalized complex traits loci identified 107 fetal- and 30 piglet-stage regulatory elements, with only three eGenes shared between stages.

These findings provide insights into developmental chromatin dynamics in skeletal muscle and an effective framework for prioritising putative regulatory variants affecting pig traits at prenatal stages.

## Introduction

Myogenesis, the process of muscle formation, is controlled by a tightly regulated network of genetic and epigenetic factors, including chromatin accessibility, transcription factors, and cis-regulatory elements [1]. In the context of the Functional Annotation of ANimal Genomes (FAANG) global project [2,3], the chromatin functional states of the skeletal muscle have been extensively annotated in various livestock species, including pigs [4,5]. However, the regulatory landscape of porcine muscle across prenatal and postnatal developmental stages remains incompletely characterised to date. Most of the available studies [6–10] have relied on the Assay for Transposase-Accessible Chromatin with sequencing (ATAC-seq) [11] to highlight the dynamic changes in chromatin accessibility occurring between various prenatal and postnatal stages. For example, Salavati et al. [7] identified more than 4,600 regions of differential accessibility in the semitendinosus muscle of fetuses at days 45, 60, and 90, as well as in piglets aged one and six weeks. Many of these regions were found to be associated with genes involved in myogenesis and muscle growth.

Population-based molecular quantitative trait locus (molQTL) mapping, statistically associating genomic variants with molecular phenotypes (for example, gene expression and epigenetic modifications), is a key approach to better understanding the biological mechanisms underlying complex phenotypes *in vivo* [12,13]. Among the resources already developed for farmed animals by the FarmGTEx project [14], the Porcine Genotype-by-Tissue-by-Expression Atlas, PigGTEx (http://piggtex.farmgtex.org) has catalogued tens of thousands of cis-eQTLs (expression Quantitative Trait loci) and eGenes (i.e., genes whose expression is regulated by at least one cis-eQTL), as well as splicing QTLs (sQTLs), exon expression QTLs (eeQTLs), enhancer QTLs (enQTLs) and long non-coding RNA QTLs (lncQTLs) across multiple pig tissues, including skeletal muscle [15].

The objective of the present study was to identify developmental stage–specific accessibility regions in the pig skeletal muscle and to highlight putative causal regulatory variants underlying complex traits. The scheme of the analytical workflow is provided in Figure 1. We first generated a comprehensive chromatin accessibility resource for porcine skeletal muscle by aggregating and reprocessing four publicly available ATAC-seq datasets spanning various prenatal and postnatal (hereafter referred to as ‘fetal’ and ‘piglet’, respectively) developmental stages. This cross-study integration allowed expanding breed diversity and developmental coverage, thereby enabling the identification of more robust and generalizable consensus peaks and DAPs (Differentially Accessible Peaks) (Fig. 1A). We then integrated both consensus peaks and DAPs with the comprehensive set of cis-acting skeletal muscle molQTLs from the PigGTEx project [15] (Fig. 1B) and applied gene set enrichment analysis (GSEA) to interpret the functional programs associated with eGenes at each stage. We further examined the DAPs at promoter–TSS (transcription start sites) overlapping the strongest eQTLs to assess how the accessibility shifts at these sites occurred in fetal and piglet muscle. In addition, we tested the enrichment and genomic distribution across fetal and post-natal stages of the strongest effect variants (top 25% of each molQTL class) within these DAPs (Fig. 1C). Finally, we incorporated the gene–trait colocalization results derived from a large-scale integrative GWAS framework developed by Xu et al. [16] (Fig. 1D). This framework combines transcriptome-wide association studies (TWAS), summary-based Mendelian randomization (SMR), and Bayesian colocalization to identify loci where gene expression and trait variation likely share a common causal variant. By intersecting these colocalized gene–trait associations with promoter-accessible eGenes, we highlighted 107 fetal-stage and 30 piglet-stage regulatory elements associated with complex traits.

**Figure 1.**
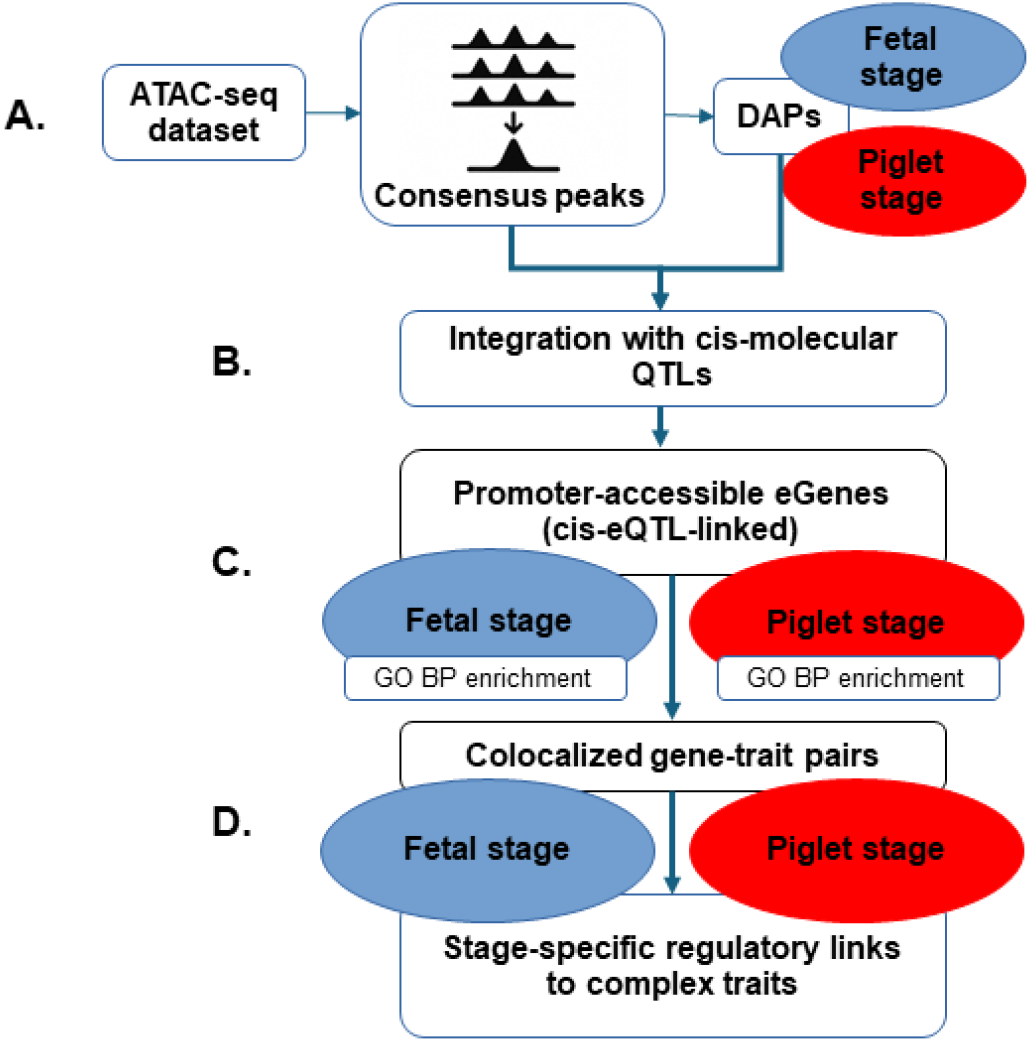
Scheme of the analytical workflow. **A)** Four publicly available ATAC-seq datasets were aggregated to generate consensus peaks, followed by genomic annotation and differential accessibility analysis to identify DAPs (Differentially Accessible Peaks) across the two developmental stages. **B)** Consensus peaks and DAPs were integrated with cis-molQTLs from the PigGTEx resource. **C)** eGenes associated to promoter eQTLs located inside DAPs were subjected to GO enrichment analysis and **D)** intersected with colocalized GWAS gene–trait pairs to identify developmental stage–specific regulatory elements linked to gene expression and complex traits in porcine skeletal muscle.

Through this multi-layered integration of functional genomic, molecular QTLs, and trait association data, our study provides new insights into the developmental dynamics of chromatin accessibility in the skeletal muscle and an effective method of prioritising putative regulatory variants that affect pig traits at prenatal stages.

## Materials and methods

### ATAC-seq dataset selection

Raw FASTQ were retrieved from four bulk ATAC-seq datasets of porcine skeletal muscle, comprising a total of 202 animals (110 fetal and 92 piglet samples; see Suppl. Table 1). All the ATAC-seq libraries were sequenced on the Illumina NovaSeq platform, minimising any potential batch effects related to the sequencing technology used, and metadata on tissue type, developmental stage, sex, and dataset source were available for each experiment. Only these four datasets met these uniform metadata criteria at the time of the search; the others were excluded due to different target tissues, inaccessible raw data, or insufficient metadata for integration.

The first two datasets were generated by the GENE-SWitCH project (https://data.faang.org/projects/GENE-SWitCH). Dataset 1 (12 Large White samples) was obtained from fetuses on days 30 and 70, and from newborn piglets (https://data.faang.org/dataset/PRJEB44468). Dataset 2 (160 samples from a cross between TN60/TN70 sows and Duroc/Tempo boars) was obtained from fetuses on day 70 and from 10-week-old piglets (https://data.faang.org/dataset/PRJEB53440). Dataset 3 (24 Large White × Landrace crossbred pigs) was generated by Salavati et al. [7] from fetuses on days 45, 60, and 90 and from one- and six-week-old piglets (https://data.faang.org/dataset/PRJEB41485). Dataset 4 (six samples from Luchuan and Duroc pigs) was generated by Miao et al. [17] from six-month-old pigs (https://www.ebi.ac.uk/ena/browser/view/PRJNA749761). Although the majority of samples were provided by dataset 2, incorporating the additional datasets increased breed diversity and the number of developmental time points, thereby reducing study-specific biases.

Uniform downstream processing was performed on all samples. All the sampling time points were retained and datasets were merged and categorized into two broad prenatal and postnatal developmental stages (‘fetal’ and ‘piglet’) due to the strong imbalance of sample sizes across time points.

### Alignment and peak calling of compiled ATAC-seq data

Raw sequencing reads from the compiled ATAC-seq datasets were processed using version 1.2.1 of the nf-core ATAC-seq pipeline [18] (https://nf-co.re/atacseq). Reads were aligned to the Sus scrofa 11.1 reference genome (Ensembl release 108; NCBI RefSeq accession GCF_000003025.6) using BWA [19], and quality control metrics were assessed within the pipeline.

For peak calling, MACS2 [20] was used to identify OCRs in each group. Peaks were first called per sample with MACS2 and then merged into a single consensus peak set across all samples using the nf-core workflow parameter (--min_reps_consensus 2) which requires a peak to be present in at least two biological replicates within a group to be retained. Annotation of consensus peaks was performed using HOMER, which assigns peaks to genes based on direct overlap with a transcription start site (TSS) or to the nearest gene when no overlap is present (default nearest-gene assignment without distance cutoff). Peak quantification was performed using Subread featureCounts to count the number of aligned reads overlapping consensus peak regions; no minimal read threshold was applied, i.e., all consensus peaks were retained at this stage. The generated count matrix was used for normalization and differential accessibility analyses. Mitochondrial peaks were removed as artifacts before downstream analyses.

### Normalization and differential accessibility analysis

To correct for non-linear trended efficiency biases, loess normalization was applied to peak counts using the csaw Bioconductor package [21]. Principal component analysis (PCA) was performed on log-normalized accessibility data to visualize genome-wide variation between the two developmental stages.

Peaks with globally weak accessibility were filtered out, retaining only those with greater than 10 counts per million (CPM) in at least 10 samples. Differential accessibility analysis was performed on log-normalized CPM values using the limma-voom approach [22]. Empirical sample quality weights were included to account for experimental biases that systematically affected variability for all or most of the peaks in a given sample. For each peak, a linear model was fitted, with sex and developmental stage as fixed factors, and experiment (dataset source) as a random factor to account for batch effects. Contrasts were estimated to test for differential accessibility between developmental stages (piglets vs fetuses) and sexes (males vs females).

Multiple testing correction was performed using the Benjamini-Hochberg procedure, with a false discovery rate (FDR) threshold of 5% to identify statistically significant DAPs.

### Integrative analysis of ATAC-seq peaks and cis-molQTLs

Annotated ATAC-seq peaks were integrated with multiple types of cis-acting molQTLs from the PigGTEx project [15]. For each type of molQTL (eQTLs, sQTLs, eeQTLs, enQTLs, lncQTLs) in skeletal muscle, cis-molQTLs were identified using a window of ±1 Mb around the TSS of the corresponding gene. Significant cis-molQTLs were defined using an FDR threshold of < 0.05, implemented as a Local False Sign Rate (LFSR) < 0.05.

To identify overlapping genomic regions between ATAC-seq peaks and cis-molQTLs, analyses were conducted using the GenomicRanges (version 1.58.0) package in R [23]. No minimum overlap threshold was applied, meaning peaks and cis-molQTLs overlapping by even a single base pair were considered to be a valid intersection. The same approach was used to identify overlaps between DAPs and cis-molQTLs.

For eQTLs specifically, promoter–TSS overlaps were examined in more detail. We ranked variants by absolute effect size and selected the top 1% strongest eQTLs. Among these, only those falling within promoter–TSS DAPs were retained. We then quantified their logFC in accessibility between stages to assess whether the strongest regulatory variants preferentially coincided with fetal or piglet promoters.

Beyond promoters, we assessed whether strong-effect variants across all molQTL classes were preferentially enriched within DAPs. Strong variants were defined as the top 25% of each molQTL class based on absolute effect size. Enrichment was tested using Fisher’s exact test, comparing strong versus non-strong variants inside versus outside DAPs. Finally, we stratified these strong variants by genomic category (intergenic, promoter–TSS, exon, intron, TTS) to evaluate their developmental distribution between fetal and piglet muscle. Promoter–TSS regions were defined using the HOMER’s default annotation criteria (−1 kb to +100 bp relative to the TSS).

### Integration with trait–gene colocalization data

We integrated our ATAC-seq results with the trait–gene co-localization data generated by Xu et al. [16]. This study combined transcriptome-wide association studies (TWAS), summary-data Mendelian randomisation (SMR) and Bayesian co-localization to identify loci where gene expression and phenotypic variation likely share a causal variant. Only gene–trait pairs with a regional co-localization probability (RCP) greater than 0.5 were retained. The trait set comprised two complementary meta-analytical designs: (i) cross-breed meta-analyses for 232 traits based on 2,056 high-quality GWAS summary statistics across 59 pig populations (labeled M_), and (ii) breed-specific meta-analyses for 12 traits in Duroc (D_), Landrace (L_), and Yorkshire (Y_) pigs, capturing lineage-specific effects [16].

We intersected the trait-associated genes identified by Xu et al. [16] with the stage-specific eGenes (i.e., genes whose expression is regulated by at least one cis-eQTL) associated with accessible promoter regions identified in our ATAC-seq analysis, enabling us to link chromatin accessibility to trait-associated gene regulation in pig skeletal muscle.

### Gene set enrichment analysis (GSEA)

We performed Gene Ontology (GO) enrichment analysis to explore the biological processes associated with promoter-accessible eGenes linked to cis-eQTLs in fetal and piglet muscle. Enrichment analysis was conducted separately for genes associated with peaks of increased accessibility in fetuses and those with increased accessibility in piglets using the DAVID Functional Annotation Tool [24]. We focused on GO Biological Process categories with uncorrected p-values < 0.05, fold enrichment > 1.5, and a minimum of three genes per term to identify potential functional trends within each developmental stage. The background gene list was set to the default background in DAVID.

## Results

### Compiled chromatin accessibility landscape in porcine skeletal muscle

To characterize the chromatin accessibility profile of porcine skeletal muscle, open chromatin regions were identified from ATAC-seq data generated across 202 muscle samples, including 110 fetuses and 92 piglets (Suppl. Table 1). The datasets were merged and grouped into two broad developmental stages: fetal and piglet. Sequencing reads from the combined dataset were aligned to the reference pig genome Sus scrofa 11.1, and OCRs were subsequently identified.

A total of 170,978 consensus peaks were identified through genome-wide chromatin profiling. Genomic annotation showed that 80,676 of these peaks were located in intronic regions, 62,225 in intergenic regions, 12,896 in exons, 10,429 in promoter-TSS regions, and 4,048 in transcription termination site regions (Suppl. Table 2). This distribution was consistent with the expected positioning of cis-regulatory elements in the pig and other mammalian genomes [25,26]. The read distribution profiles around transcription start sites (TSS) showed a strong enrichment at the TSS and upstream regions (not shown), consistent with expected patterns of promoter-associated chromatin accessibility [9].

PCA was performed on the normalized chromatin accessibility profiles based on consensus peaks to explore the global variation across samples (Figure 2). The PCA revealed mild stratification according to the dataset origin, indicating small but detectable batch-associated effects across experiments. As expected, and as shown in previous work [9], no clear separation was observed between male and female samples (not shown). In contrast, fetal and piglet samples were clearly separated along the first principal component, confirming the widespread developmental differences in chromatin accessibility patterns previously observed [9].

**Figure 2.**
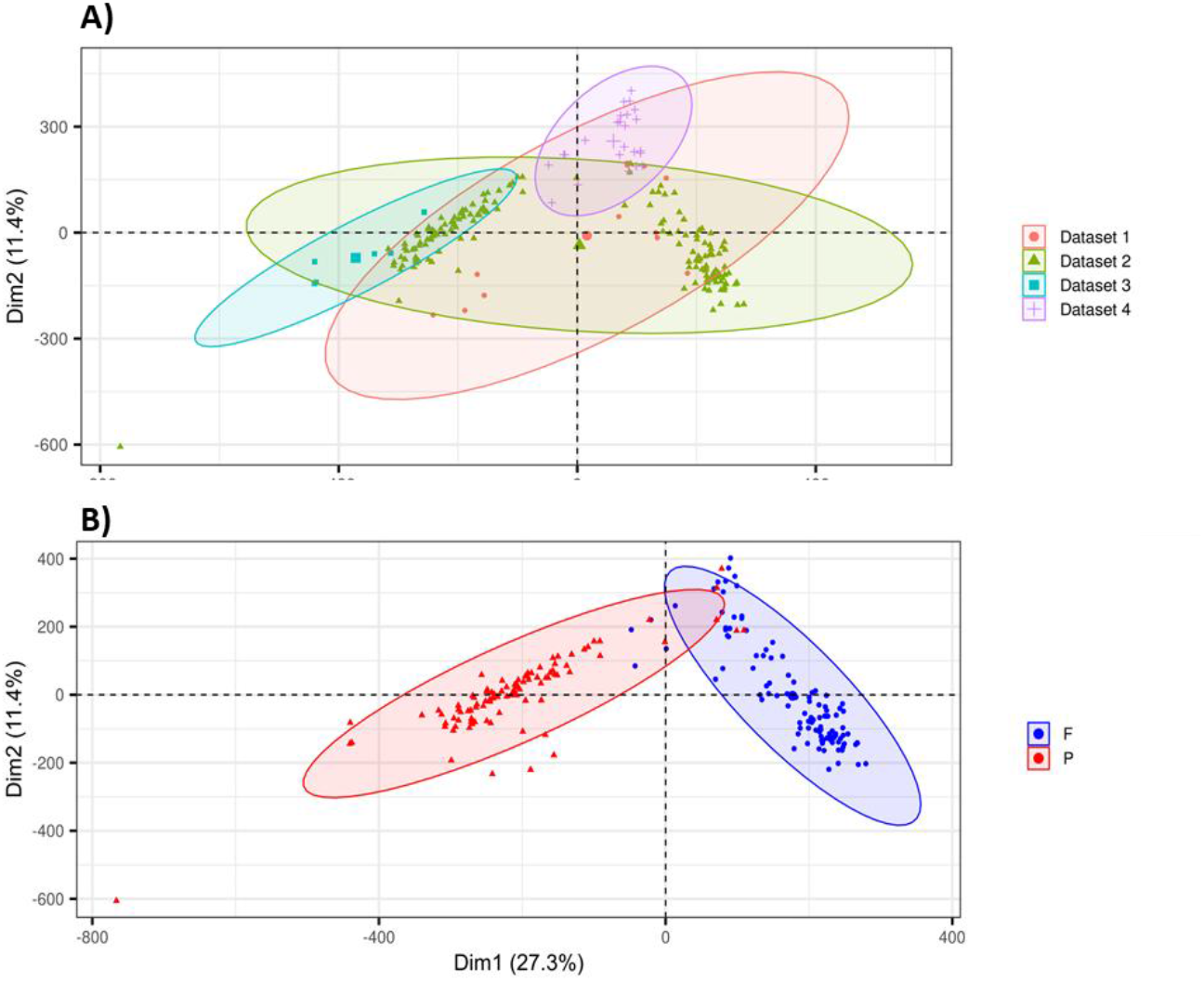
Principal component analysis (PCA) of LOESS-normalized chromatin accessibility profiles based on consensus ATAC-seq peaks. Each point represents a sample, and ellipses summarize group-wise variation across principal components. Samples are colored by A) experiment of origin where PCA shows a mild batch-associated effects across different studies and B) developmental stage where PCA shows a clear developmental separation between fetal and piglet samples along the first principal component.

### Differential chromatin accessibility between fetal and piglet stages

The differential accessibility analysis was conducted between fetal and piglet samples using the limma-voom framework with empirical quality weights. After excluding peaks on sex chromosomes, a total of 132,275 significant DAPs between developmental stages were identified at an FDR threshold of 0.05. Among these, 70,756 peaks exhibited increased accessibility in fetal muscle, while 61,519 were more accessible in piglet muscle. Each differentially accessible peak was thus assigned to the developmental stage in which it was more accessible. Note that this classification reflected statistically significant differences in accessibility, rather than mere presence or absence of signal.

Genomic annotation revealed that DAPs followed a distribution similar to the consensus peaks, with the majority located in intronic and intergenic regions. However, stage-specific differences in chromatin accessibility were evident (Figure 3; Suppl. Table 3). Significant differences were found in the distribution of DAPs between developmental stages (χ^2^ test, *P* < 2.2 × 10^−16^). Peaks more accessible in fetal muscle were enriched in intergenic regions (28,654 vs. 18,764; Fisher’s exact test, *P* = 1.65 × 10^−314^) and promoter–TSS regions (5,786 vs. 2,556; *P* = 4.52 × 10^−204^), whereas piglet-associated peaks were enriched in intronic regions (34,239 vs. 29,525; *P* < 2.2 × 10^−16^).

**Figure 3.**
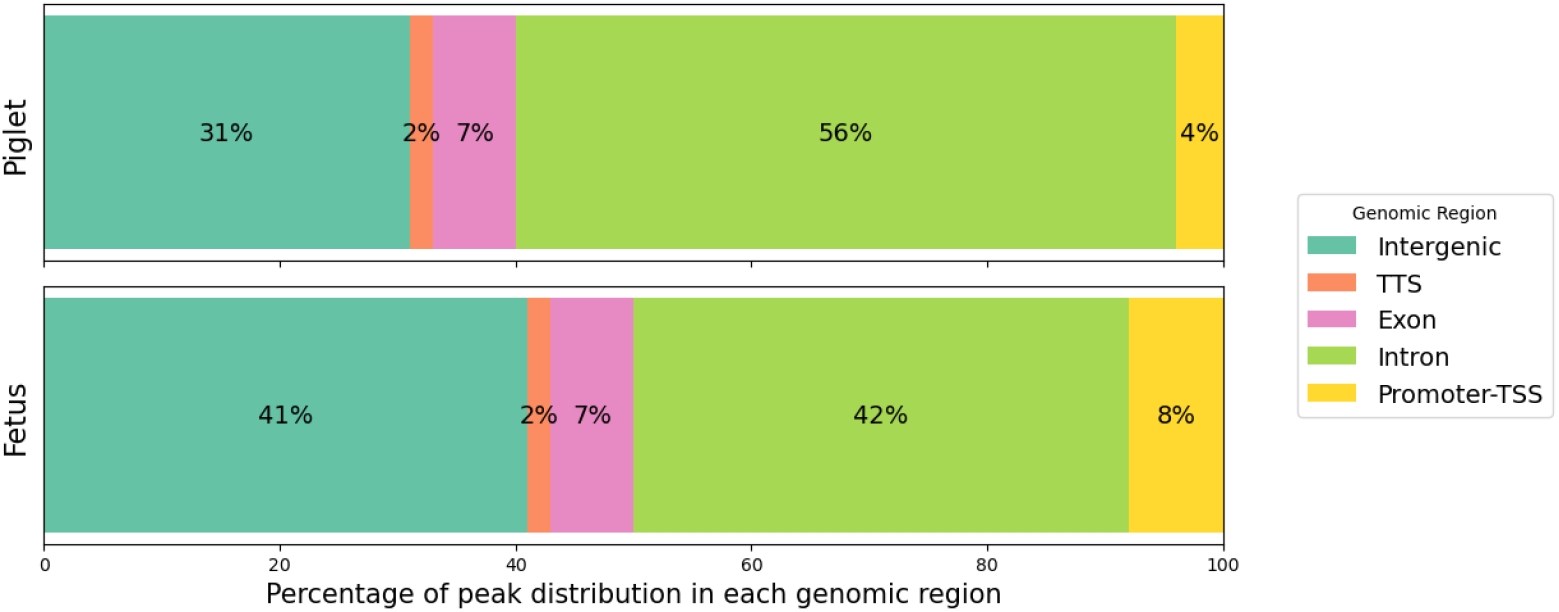
Genomic distribution of differentially accessible peaks (DAPs) between fetal and piglet skeletal muscle. Stacked bar plots show the percentage of DAPs located in intergenic, intronic, exonic, promoter–TSS, and transcription termination site (TTS) regions for each developmental stage.

### Integration of chromatin accessibility and molecular QTLs

#### -Integration of ATAC-seq consensus peaks with molQTLs

To investigate the regulatory potential of accessible chromatin in porcine skeletal muscle, we explored the overlap between chromatin accessibility and the PigGTEx dataset of cis-acting molQTLs [15]. Supplementary Table 4 provides a summary of the skeletal muscle cis-molQTLs, the total number of variants and their overlaps with consensus ATAC-seq peaks and DAPs between the fetal and piglet stages. Muscle-specific cis-molQTLs were identified by associating genetic variants with proximal molecular traits to identify context-dependent regulatory variants that mechanistically link genotype to transcriptional regulation.

We first intersected molQTLs and consensus ATAC-seq peaks (Figure 4, Suppl. Table 5A). Among the 162,807 consensus peaks, 60,554 (37.2%) overlapped with 343,496 cis-eQTL variants, corresponding to 9,093 unique eGenes (Suppl. Table 6). The distribution of cis-eQTL-overlapping peaks roughly reflected that of genomic annotation regions: intronic (19.4%) and intergenic (9.0%) were most frequent, followed by exonic (3.9%), promoter-TSS (3.7%), and TTS (1.4%) regions. A total of 50,150 (30.8%) consensus peaks overlapped with 275,864 sQTL variants, with similar genomic distributions: intronic (16.4%), intergenic (7.2%), exonic (3.4%), promoter-TSS (3.3%), and TTS (1.2%) peaks. Enhancer QTLs (enQTLs) were found in 23,530 (14.5%) consensus peaks, overlapping 107,021 variants, primarily in intronic (7.3%) and intergenic (3.2%) regions. For exon expression QTLs (eeQTLs), 58,744 (36.1%) consensus peaks overlapped with 327,363 variants, again enriched in intronic (18.8%) and intergenic (8.4%) contexts. Lastly, 34,944 (21.5%) consensus peaks overlapped with 160,339 lncQTL variants, with the highest representation in intronic (11.0%) and intergenic (5.0%) regions (Suppl. Table 5A).

**Figure 4.**
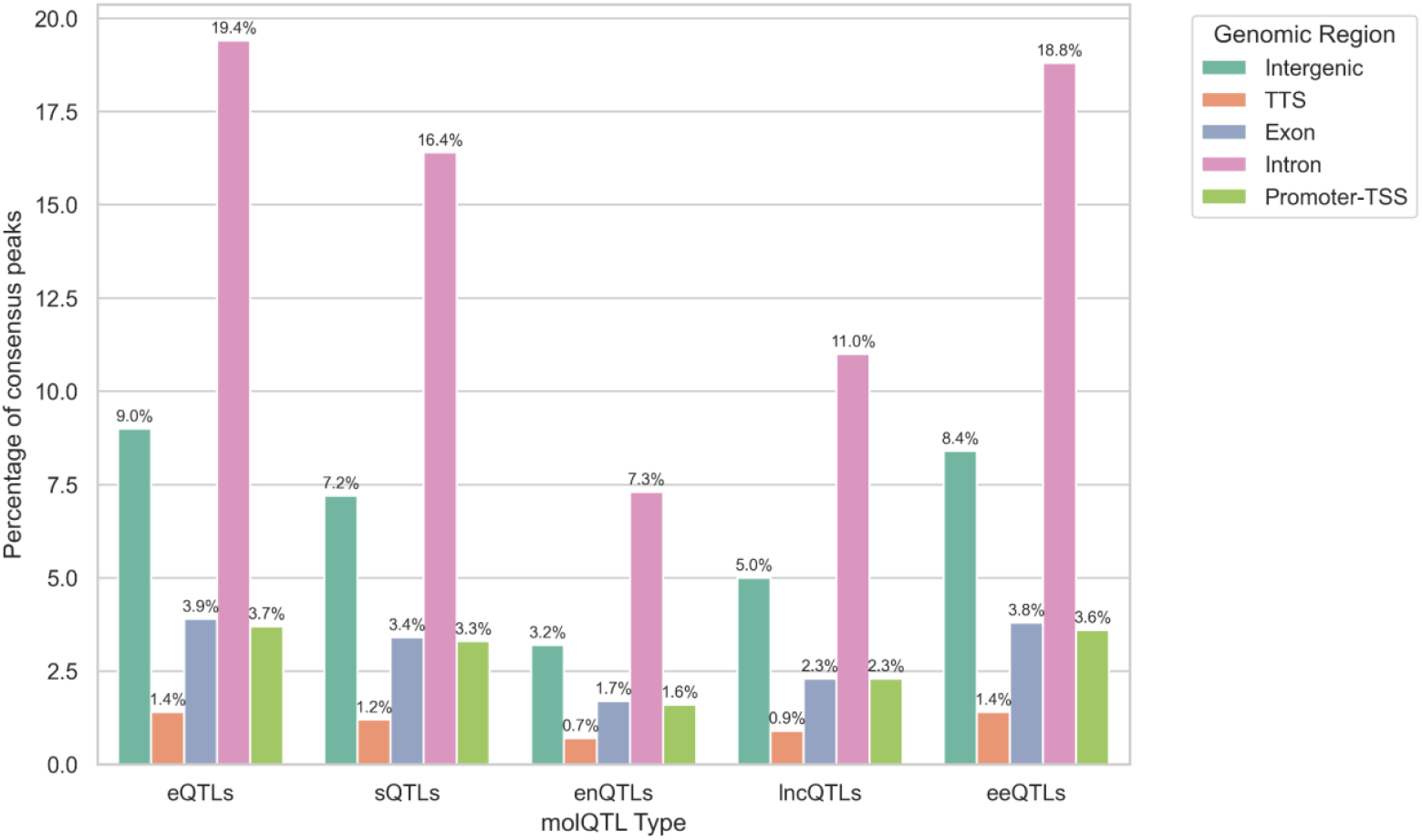
Genomic annotation of consensus ATAC-seq peaks overlapping with cis-regulatory molQTLs in porcine skeletal muscle. Bars represent the percentage of consensus peaks overlapping with eQTLs, sQTLs, enQTLs, lncQTLs, and eeQTLs, subdivided by genomic region. For all molQTL types, the number of overlapping variants was significantly greater than expected by chance based on 1,000 permutations (empirical *P* < 0.001).

To statistically assess whether the observed overlaps exceeded random expectation, we performed a permutation-based enrichment analysis. For each molQTL class, we generated 1,000 randomized sets of peak coordinates preserving peak count, length, and chromosomal assignment. The number of overlapping molQTL variants was computed for each permutation to derive a null distribution. For all the eQTLs (343,496), sQTLs (275,864), eeQTLs (327,363), enQTLs (107,021), and lncQTLs (160,339),the observed number of overlapping variants was significantly greater than expected by chance (empirical *P* < 0.001).

#### -Integration of DAPs between fetal and piglet stages with molQTLs

To assess how developmental changes in chromatin accessibility are associated with regulatory variants, we next examined the overlap of DAPs and molQTLs at the fetal and piglet stages.

A total of 25,008 (15.4%) DAPs with higher accessibility in fetuses and 25,725 (15.8%) DAPs with higher accessibility in piglets overlapped with at least one cis-eQTL region, corresponding to 153,325 and 141,374 eQTLs (Suppl. Table 5B) and 8,266 (Suppl. Table 7) and 8,177 eGenes (Suppl. Table 8), respectively. Although the total number of overlapping peaks was comparable between developmental stages, promoter/TSS-associated DAPs exhibited a notable asymmetry, both in the percentage of DAPs overlapping eQTLs (2.2% in fetus vs. 1.0% in piglet) and in the number of associated genes (1,948 vs. 716) that were both annotated by HOMER and classified as eGenes, indicating a stronger regulatory signal in promoter regions during fetal muscle development (Figure 5).

**Figure 5.**
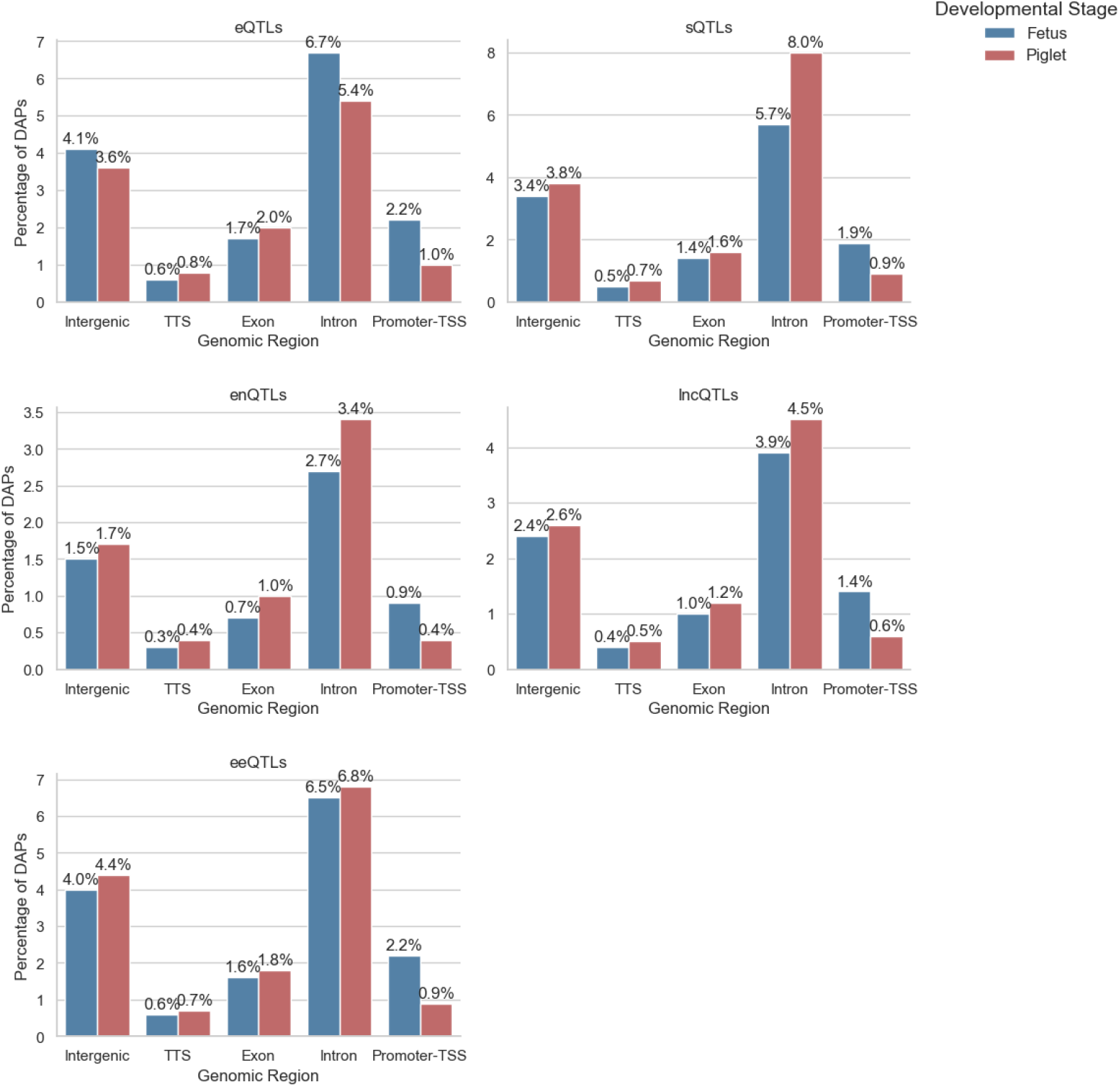
Stage-specific distribution of differentially accessible peaks (DAPs) overlapping molQTLs. Bar plots show the percentage of fetal and piglet-accessible DAPs that overlap molQTLs, subdivided by genomic annotation and stratified by molQTL type (eQTL, sQTL, eeQTL, enQTL, lncQTL). For all molQTL types and both developmental stages, the number of overlapping variants was significantly greater than expected by chance based on 1,000 permutations (empirical *P* < 0.001).

Overlap patterns for sQTLs followed a similar trend. Among DAPs, 21,146 (13.0%) fetal-accessible and 21,750 (13.4%) piglet-accessible peaks overlapped with 120,160 and 109,868 sQTL variants, respectively. Intronic regions were again the most frequently represented, with 5.7% fetal and 8.0% piglet intronic peaks overlapping sQTLs. Promoter-TSS associated sQTL overlaps were more frequent in fetal peaks (1.9%) than in piglet peaks (0.9%).

The eeQTLs were found in 24,261 (14.9%) fetal-accessible and 25,001 (15.4%) piglet-accessible peaks, overlapping with 145,795 and 135,630 variants, respectively. Again, introns and intergenic regions were predominant, while 3,500 (2.2%) fetal and 1,537 (0.9%) piglet promoter peaks overlapped with eeQTLs.

The integration with lncQTLs identified 14,731 (9.1%) fetal-accessible and 14,560 (8.9%) piglet-accessible peaks overlapping with 72,808 and 64,702 lncQTL variants, respectively. Intronic and intergenic annotations remained the most common, while 2,196 (1.4%) fetal versus 947 (0.6%) piglet promoter-associated peaks overlapped with lncQTLs.

Finally, in the case of enQTLs, 10,101 (6.2%) fetal-accessible peaks and 9,543 (5.9%) piglet-accessible peaks overlapped with 49,518 and 42,394 enQTL variants, respectively. The majority of these were intronic (4,392 in fetus; 5,502 in piglet), followed by intergenic, exonic, and promoter/TSS-associated regions. Promoter-overlapping peaks showed 1,523 (0.9%) overlaps in fetal samples versus 690 (0.4%) in piglets.

Across all molQTL types, permutation-based enrichment tests confirmed that the observed number of variant overlaps in both fetal and piglet-accessible peaks was significantly greater than expected by chance (empirical *P* < 0.001).

These results showed that chromatin regions with developmental stage-specific accessibility were significantly enriched in multiple classes of cis-regulatory variants. The annotation of these regions also suggested differential regulatory engagement between stages: fetal-accessible regions showed higher counts in promoter and intergenic compartments, while piglet-accessible regions were more represented in intronic contexts.

#### -Enrichment of strong-effect molQTL variants in differentially accessible peaks

Next, we asked whether chromatin accessibility changes preferentially coincide with strong-effect molQTLs across the five classes of molecular QTLs tested. For each dataset, the strongest 25% of variants were defined based on absolute effect size. Although only a minority of the strong molQTLs overlapped DAPs (∼6–9.5%, depending on molQTL type), a substantial fraction of the molQTLs present within DAPs were strong-effect variants (∼32–52%). Enrichment analysis confirmed that strong molQTLs were significantly more likely than expected to coincide with DAPs in both fetal and piglet muscle (odds ratios 1.1–1.2, *P* = 1 × 10^−67^–1 × 10^−235^; Suppl. Table 9). These results indicate that accessibility differences between developmental stages harbour strong-effect regulatory variants across multiple molecular QTL types.

Then we examined the genomic distribution of these strong-effect variants within DAPs (Figure 6). Intronic regions accounted for the majority of overlaps across all molQTL types, with ∼43–57% of strong variants and a consistent piglet bias (∼56% vs ∼43% in fetus). Promoter–TSS regions, although representing a smaller proportion (∼8–17%), showed a pronounced fetal bias, with significantly more strong-effect variants in fetuses than in piglets. Additional differences were also detected in intergenic, exon, and TTS categories for selected molQTL classes. These results highlight how the strongest cis-regulatory events of the genome shift during development, from gene-distal and promoter-linked control at the fetal stage towards intragenic regulation at the piglet stage.

**Figure 6.**
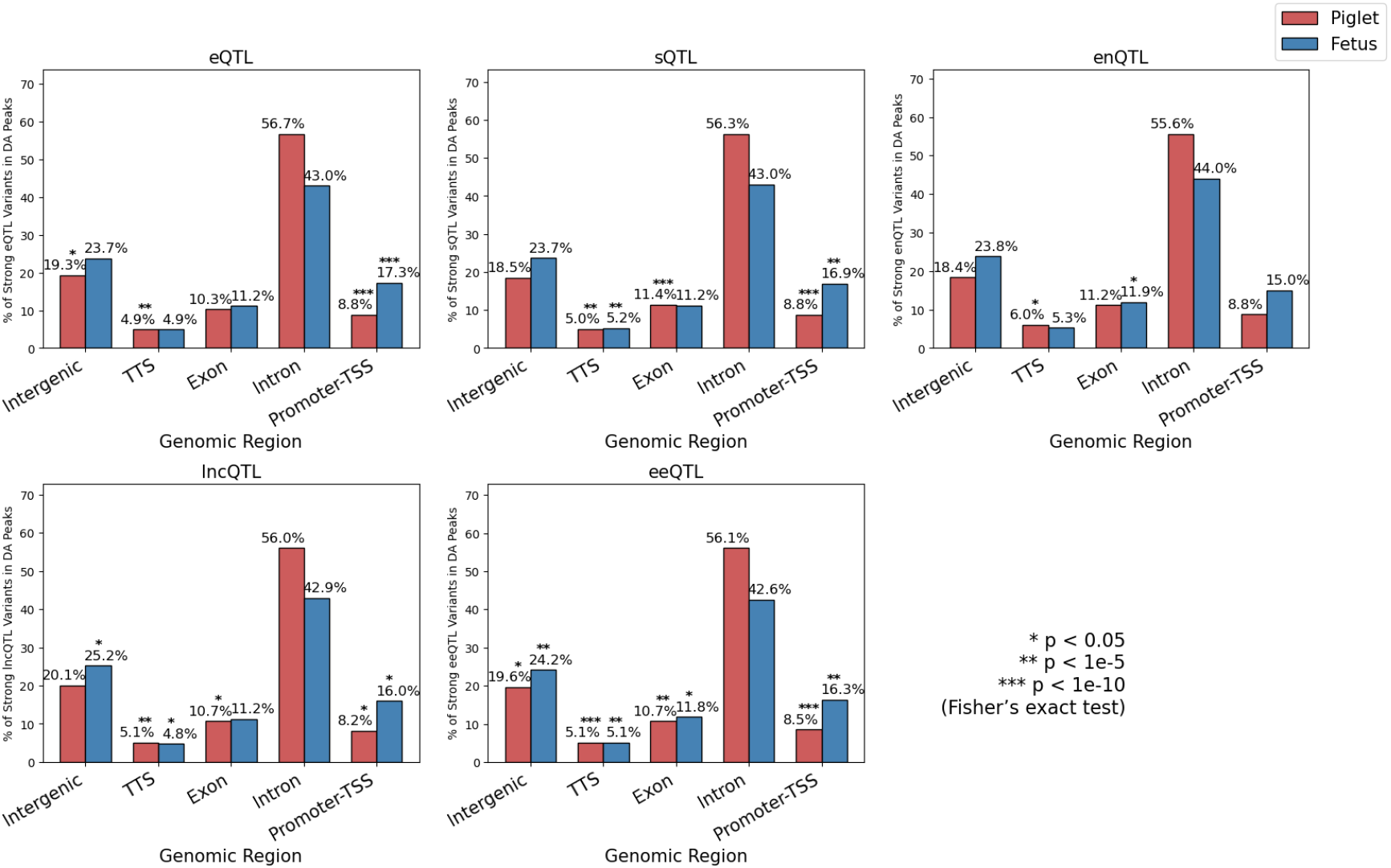
Genomic distribution of strong-effect molQTL variants within differentially accessible peaks. Barplots show the percentage of strong variants (top 25% by absolute effect size) for each molQTL type: eQTL, sQTL, enQTL, lncQTL, and eeQTL, found within differentially accessible (DA) chromatin regions in fetal (blue) and piglet (red) muscle. Genomic categories include intergenic, transcription termination site (TTS), exon, intron, and promoter–TSS regions. Asterisks indicate significant enrichment differences between fetal and piglet within each category based on Fisher’s exact test (*P* < 0.05, *P* < 1e–5, *P* < 1e–10).

We finally focused on the top 1% strong eQTLs ranked by effect size overlapping promoter–TSS DAPs. As shown in Figure 7, only four variants were located in promoter peaks more accessible in piglets, while the majority of variants (14 out of 18) overlapped promoter peaks more accessible in fetuses.

**Figure 7.**
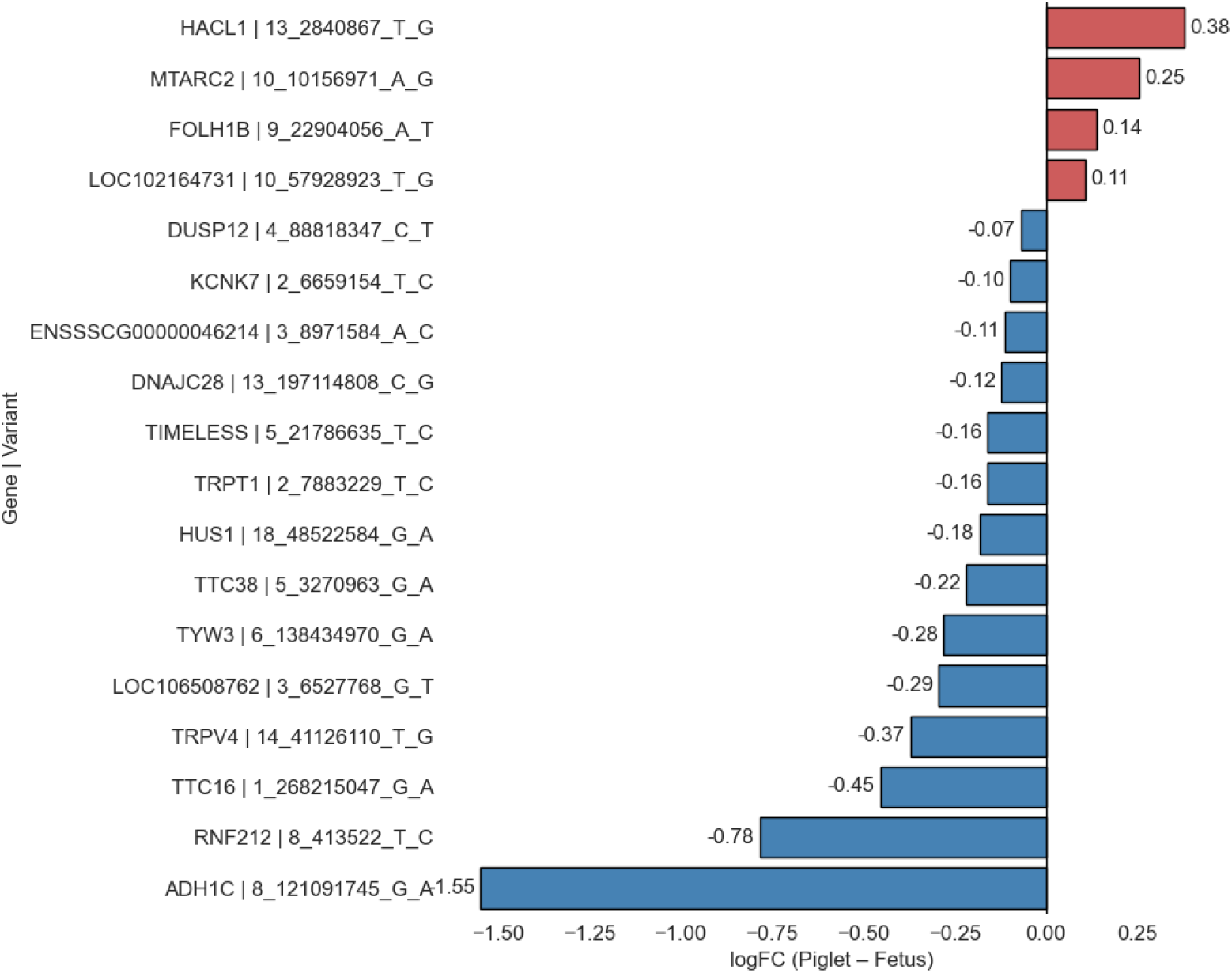
Accessibility bias of promoter–TSS eQTLs with the strongest effect. Barplots show logFC (Piglet vs. Fetus) in promoter accessibility for the top 1% strong-effect eQTLs. Four variants overlapped promoter peaks more accessible in piglets (in red), while 14 overlapped peaks more accessible in fetuses (in blue), indicating a predominant fetal bias.

### Exploratory GO term enrichment of promoter associated eGenes

To investigate the potential biological functions of genes associated with promoter-accessible regions linked to cis-eQTLs, we performed a Gene Ontology (GO) enrichment analysis separately for the fetal- and piglet-accessible gene sets. We focused in particular on GO Biological Process categories with uncorrected *P* < 0.05, fold enrichment > 1.5, and minimum gene counts ≥3 to identify possible functional trends.

Genes associated with fetal-accessible peaks linked to cis-eQTLs (n = 1,948) were enriched for RNA metabolic processes, chromosome organization, DNA repair, and ribosome biogenesis, reflecting active transcriptional and biosynthetic programs likely supporting cell proliferation and differentiation during prenatal muscle development. In contrast, piglet-accessible genes linked to cis-eQTLs (n = 716) were enriched for striated muscle contraction, lipid modification, cellular respiration, and musculoskeletal movement, consistent with the metabolic and contractile functions of postnatal skeletal muscle (Suppl. Table 10).

Notably, most terms were unique to only one developmental stage. For instance, musculoskeletal movement and striated muscle contraction were highly enriched in piglet-specific genes alone, whereas the metabolic and biosynthetic processes for nucleic acids terms were observed exclusively in the fetal set. These differences highlight how regulatory accessibility at promoters mark distinct functional states aligned with developmental transitions in muscle (Figure 8).

**Figure 8.**
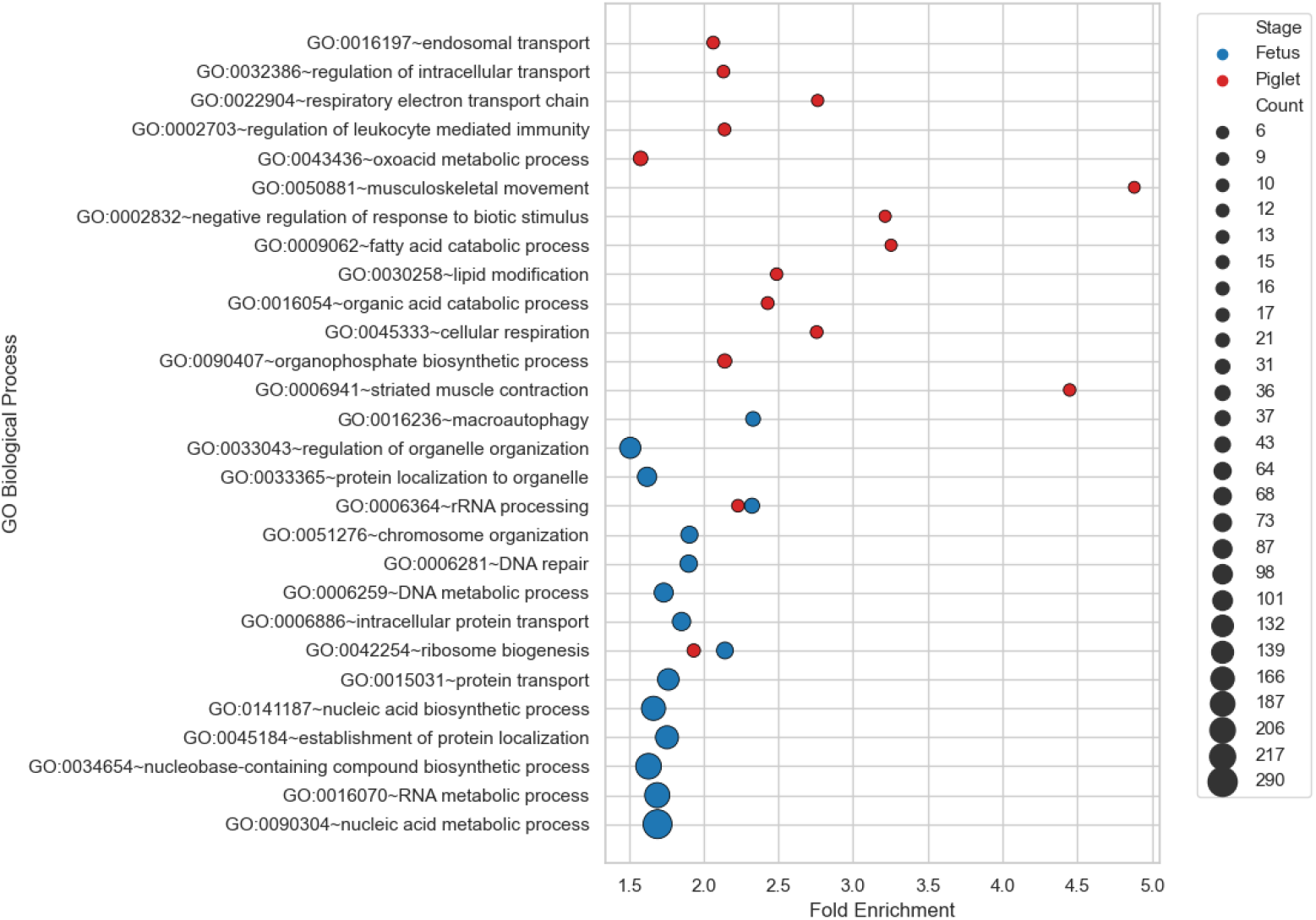
Exploratory GO term enrichment of promoter-associated eGenes in fetal and piglet skeletal muscle. Top 15 GO Biological Process (BP) terms per developmental stage enriched among genes linked to promoter-accessible, cis-eQTL-overlapping peaks. Dot size reflects gene count; X-axis shows fold enrichment. Terms shown passed filters of *P* < 0.05, fold enrichment > 1.5, and ≥ 3 genes.

We used the colocalized gene–trait associations from Xu et al. [16], restricted to pairs with RCP > 0.5, to determine which complex traits were linked to either fetal- or piglet-stage increased chromatin accessibility. Specifically, we intersected the colocalized gene–trait pairs with our dataset of promoter-associated eGenes to identify which of the genes implicated in trait variation showed developmental stage–specific chromatin accessibility at their promoters.

We identified 107 fetal-stage and 30 piglet-stage promoter-accessible eGenes colocalized with the same set of 10 complex traits (Suppl. Table 11) including growth-related (backfat thickness, BFT; average daily gain, ADG; days to reach weight, DAYS), muscle structure (loin muscle area, LMA; loin muscle depth, LMDEP; lean cut percentage, LEANCUTP), and reproduction (number born alive, NBA; number born healthy, NBH; total number born, TNB; total litter weight at birth, TLWT_BA). Despite convergence at the trait level, the overlap of fetal and piglet genes was limited. Only three eGenes were shared, namely KYAT1 (kynurenine aminotransferase 1), AGO2 (argonaute RISC catalytic component 2) and KIAA1191 (a gene to enable oxidoreductase activity). Thus, while both developmental stages were connected to the same set of complex traits, the underlying gene sets associated with promoter accessibility were largely distinct between them.

Figure 9 further illustrates how trait associations distribute across developmental stages and GWAS data from different breeds. As expected, cross-breed analyses (M_) captured a broad set of growth, muscle, and reproduction traits consistently detected at both fetal and piglet stages, while within-breed analyses highlighted lineage-specific associations (e.g., Duroc for BFT at both stages, Landrace for NBA/TNB in fetus, and Yorkshire for TLWT_BA at both stages). The Yorkshire breed showed a relatively high overlap of traits between stages, compared to more stage-restricted signals in Duroc and Landrace.

**Figure 9.**
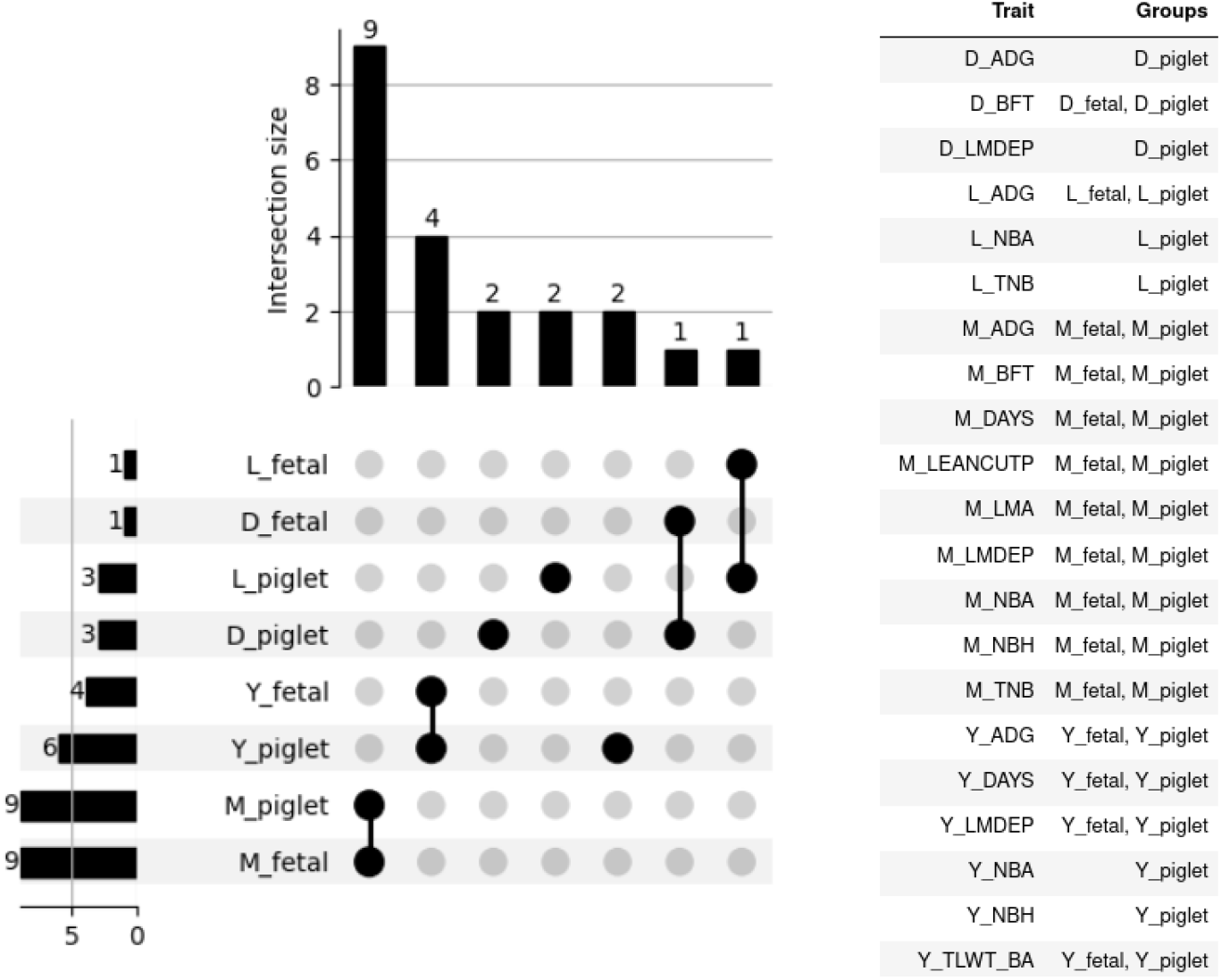
Intersection of colocalized trait associations with promoter-accessible eGenes across developmental stages and pig populations. UpSet plot showing the number and identity of traits associated with promoter-accessible eGenes colocalized with GWAS loci in fetal and piglet muscle. Traits are grouped by developmental stage (Fetus or Piglet) and by GWAS population label. The M_ prefix refers to traits derived from cross-breed meta-analyses (integrating data across 59 pig populations), while D_, L_, and Y_ indicate within-breed meta-analyses in Duroc, Landrace, and Yorkshire pigs, respectively. The plot highlights the distribution of shared and stage-specific trait associations.

## Discussion

By integrating high-quality ATAC-seq datasets across developmental stages and from different breeds, we generated a robust and generalizable map of the muscle chromatin accessibility landscape, which served as the foundation for downstream analyses of differential accessibility and regulatory variant integration using the PigGTEx resource.

Our differential accessibility analysis revealed extensive remodelling of the porcine muscle regulatory landscape between fetal and postnatal stages, with over 130,000 peaks showing significant developmental differences. The observation that fetal muscle exhibits greater accessibility at promoter and intergenic regions likely reflects the broad transcriptional priming and enhancer activity required to support active myogenesis. In contrast, the enrichment of intronic accessibility in piglet muscle suggests a shift toward regulatory refinement within gene bodies that accompanies fibre hypertrophy and metabolic specialization after birth (Figure 3). These patterns are consistent with previous studies performed in cattle [27], mouse [28] and yak [29]. Previous studies in pigs have reported dynamic stage-specific accessibility. Yue et al. [6] and Hou et al. [10] showed that fetal muscle is enriched for promoter-proximal and intergenic open chromatin, supporting developmental gene regulation, while Feng et al. [8] and Tan et al. [30] demonstrated that postnatal muscle has fewer accessible peaks overall and a relative enrichment in intronic regions, in line with functional specialization of mature muscle.

The fact that a considerable proportion of consensus peaks overlap with all molQTL classes, with more than one-third overlapping with eQTLs, confirms that chromatin accessibility provides a useful proxy for identifying functional noncoding variants (Figure 4). Previous studies in cattle [31], mice [32], and chickens [33] have reported similar findings, where substantial fractions of eQTLs and other regulatory variants are enriched within open chromatin regions identified by ATAC-seq or DNase-seq, reinforcing the utility of chromatin accessibility as a cross-species marker of functional regulatory variation. The predominance of intronic and intergenic overlap highlights that the majority of functional regulatory variants act outside of proximal promoters, consistent with enhancer-mediated and distal regulation. Similar enrichments of molQTLs in distal elements have been reported in pigs [15] and other species like cattle [34] and chickens [35] across multiple tissues supporting the notion that molecular trait variation is frequently mediated by open chromatin. Importantly, our permutation tests confirmed that the overlap of molQTLs with accessible chromatin was highly significant across all classes (empirical *P* < 0.001), indicating that these co-localizations are unlikely to arise by chance and that open chromatin is a consistent hotspot for functional regulatory variants.

The integration of molQTLs with the differentially accessible chromatin regions highlighted how cis-regulatory variants engage distinct regulatory elements across development (Figure 5, Figure 6). Although the overall number of molQTL overlaps was similar between fetal- and piglet-accessible peaks, molQTL in fetal regions were enriched in in promoter and intergenic categories, whereas piglet regions were enriched in introns. This pattern is in full agreement with the functional signatures of prenatal transcriptional plasticity and postnatal metabolic specialization previously reported in porcine transcriptomic and chromatin studies [6,8,10].

Promoter-associated eGenes were preferentially involved in RNA metabolism, DNA repair, and chromosomal organization, processes that align with the high proliferative and transcriptional activity required during myogenesis. In contrast, piglet-associated genes were related to striated muscle contraction, lipid metabolism, and cellular respiration, consistent with the metabolic and contractile demands of postnatal skeletal muscle. The presence of stage-specific terms, such as chromosome organization in fetal samples and striated muscle contraction in piglets, underscored how promoter accessibility may demarcate developmental transitions in muscle biology (Figure 8). When focusing on the top 1% of strong-effect eQTLs located in promoter–TSS DAPs (Figure 7), we observed that most of them (14 out of 18) coincided with promoters more accessible in fetal muscle. This pronounced fetal bias suggests that high-effect eQTLs are preferentially embedded in regulatory elements active during fetal muscle development, possibly reflecting the higher transcriptional plasticity of the developing tissue.

The finding that both fetal and piglet promoter-accessible eGenes colocalize with the same set of 12 complex traits indicated that developmental stages converge on common phenotypic outcomes related to growth and muscle structure, in addition to participating to the phenotypic outcomes of other physiological functions (i.e., blood parameters and reproduction). However, the observation that only three eGenes were shared between the two stages highlights how these associations are mediated by distinct genes. This reinforces the idea that accessibility changes at promoters can shift which genes influence key traits without altering the overall functional domains under selection. Moreover, trait associations were unevenly distributed across developmental stages and GWAS data from different breeds (Figure 9), likely reflecting the effect of different genetic architectures across breeds [36]. Cross-breed meta-analyses integrate diverse haplotype backgrounds, potentially revealing a wider range of variants and pathways shared across development, while within-breed analyses may emphasize more homogeneous, lineage-specific signals.

The development of skeletal muscle plays a crucial role in pig production, influencing meat yield, carcass quality, and overall commercial value [37]. The finding that the vast majority of high-effect eQTLs map to promoters that are more accessible in the fetal stage indicate that analysing chromatin accessibility is an effective first step toward the identification of regulatory variants that act at the prenatal stage to define these phenotypes. Enriching the spectrum of functional annotations covering fetal stages is thus the next step in assessing the phenotypic impact of non-coding variants in the pig genome [38] and streamlining the identification of causal regulatory variants affecting complex traits at different developmental stages.

## Acknowledgements

ES thanks the Agricultural University of Tirana for providing the opportunity to undertake the research that led to this work.

## Author contributions

ES performed all the data collection and analyses, and wrote the first draft of the manuscript. AR, SD and VP supervised data integration and statistical analyses. EC and JE provided assistance with local data curation and analysis. SD co-supervised ES’s work with EG. All authors contributed to manuscript editing. EG conceived the study, supervised data interpretation and finalised the manuscript.

## Data availability statement

This study relied on publicly available ATAC-seq datasets with BioProject accession codes PRJEB44468$, PRJEB53440, PRJEB41485 and PRJNA749761. Processed data and the full summary statistics of molQTL mapping are available at http://piggtex.farmgtex.org/.

## Additional information

The authors declare no competing interests.

## References

1. Lin, H. et al. Reprogramming of cis-regulatory networks during skeletal muscle atrophy in male mice. Nat. Commun. 14, 6581 (2023).

2. Giuffra, E., Tuggle, C. K., & FAANG Consortium. Functional Annotation of Animal Genomes (FAANG): Current Achievements and Roadmap. Annu. Rev. Anim. Biosci. 7, 65–88 (2019).

3. Clark, E. L. et al. From FAANG to fork: application of highly annotated genomes to improve farmed animal production. Genome Biol. 21, 285 (2020).

4. Pan, Z. et al. Pig genome functional annotation enhances the biological interpretation of complex traits and human disease. Nat. Commun. 12, 5848 (2021).

5. Zhao, Y. et al. A compendium and comparative epigenomics analysis of cis-regulatory elements in the pig genome. Nat. Commun. 12, 2217 (2021).

6. Yue, J. et al. The landscape of chromatin accessibility in skeletal muscle during embryonic development in pigs. J. Anim. Sci. Biotechnol. 12, 56 (2021).

7. Salavati, M. et al. Profiling of open chromatin in developing pig (Sus scrofa) muscle to identify regulatory regions. G3 Bethesda Md 12, jkab424 (2022).

8. Feng, L. et al. The Landscape of Accessible Chromatin and Developmental Transcriptome Maps Reveal a Genetic Mechanism of Skeletal Muscle Development in Pigs. Int. J. Mol. Sci. 24, 6413 (2023).

9. Chalabi, S. et al. Differences in maternal diet fiber content influence patterns of gene expression and chromatin accessibility in fetuses and piglets. Genomics 117, 110995 (2025).

10. Hou, X. et al. Dynamic changes in chromatin accessibility and gene expression involved in fetal myogenesis of Min pigs. Anim. Biosci. 38, 2525–2536 (2025).

11. Buenrostro, J. D., Giresi, P. G., Zaba, L. C., Chang, H. Y. & Greenleaf, W. J. Transposition of native chromatin for fast and sensitive epigenomic profiling of open chromatin, DNA-binding proteins and nucleosome position. Nat. Methods 10, 1213–1218 (2013).

12. Aguet, F. et al. Molecular quantitative trait loci. Nat. Rev. Methods Primer 3, 4 (2023).

13. Aguet, F. et al. Genetic effects on gene expression across human tissues. Nature 550, 204–213 (2017).

14. Fang, L. et al. The Farm Animal Genotype-Tissue Expression (FarmGTEx) Project. Nat. Genet.57, 786–796 (2025).

15. Teng, J. et al. A compendium of genetic regulatory effects across pig tissues. Nat. Genet. 56, 112–123 (2024).

16. Xu, Z. et al. Integrating large-scale meta-GWAS and PigGTEx resources to decipher the genetic basis of 232 complex traits in pigs. Natl. Sci. Rev. 12, nwaf048 (2025).

17. Miao, W. et al. Integrative ATAC-seq and RNA-seq Analysis of the Longissimus Muscle of Luchuan and Duroc Pigs. Front. Nutr. 8, 742672 (2021).

18. Ewels, P. A. et al. The nf-core framework for community-curated bioinformatics pipelines. Nat. Biotechnol. 38, 276–278 (2020).

19. Li, H. & Durbin, R. Fast and accurate short read alignment with Burrows-Wheeler transform. Bioinforma. Oxf. Engl. 25, 1754–1760 (2009).

20. Zhang, Y. et al. Model-based analysis of ChIP-Seq (MACS). Genome Biol. 9, R137 (2008).

21. Lun, A. T. L. & Smyth, G. K. csaw: a Bioconductor package for differential binding analysis of ChIP-seq data using sliding windows. Nucleic Acids Res. 44, e45 (2016).

22. Liu, R. et al. Why weight? Modelling sample and observational level variability improves power in RNA-seq analyses. Nucleic Acids Res. 43, e97 (2015).

23. Lawrence, M. et al. Software for computing and annotating genomic ranges. PLoS Comput. Biol. 9, e1003118 (2013).

24. Huang, D. W. et al. The DAVID Gene Functional Classification Tool: a novel biological module-centric algorithm to functionally analyze large gene lists. Genome Biol. 8, R183 (2007).

25. Bai, J. et al. Profiling of Chromatin Accessibility in Pigs across Multiple Tissues and Developmental Stages. Int. J. Mol. Sci. 24, 11076 (2023).

26. Alexandre, P. A. et al. Chromatin accessibility and regulatory vocabulary across indicine cattle tissues. Genome Biol. 22, 273 (2021).

27. Cao, X. et al. Comparative Enhancer Map of Cattle Muscle Genome Annotated by ATAC-Seq. Front. Vet. Sci. 8, 782409 (2021).

28. Dos Santos, M. et al. Opposing gene regulatory programs governing myofiber development and maturation revealed at single nucleus resolution. Nat. Commun. 14, 4333 (2023).

29. Li, J. et al. Integration of ATAC-Seq and RNA-Seq Analysis to Identify Key Genes in the Longissimus Dorsi Muscle Development of the Tianzhu White Yak. Int. J. Mol. Sci. 25, 158 (2023).

30. Tan, W. et al. Integrative Analysis of ATAC-Seq and RNA-Seq Identifies Key Genes Affecting Muscle Development in Ningxiang Pigs. Int. J. Mol. Sci. 26, 2634 (2025).

31. Yuan, C. et al. An organism-wide ATAC-seq peak catalog for the bovine and its use to identify regulatory variants. Genome Res. 33, 1848–1864 (2023).

32. Keele, G. R. et al. Integrative QTL analysis of gene expression and chromatin accessibility identifies multi-tissue patterns of genetic regulation. PLoS Genet. 16, e1008537 (2020).

33. Zhu, X.-N. et al. Chicken chromatin accessibility atlas accelerates epigenetic annotation of birds and gene fine-mapping associated with growth traits. Zool. Res. 44, 53–62 (2023).

34. Liu, S. et al. A multi-tissue atlas of regulatory variants in cattle. Nat. Genet. 54, 1438–1447 (2022).

35. Pan, Z. et al. An atlas of regulatory elements in chicken: A resource for chicken genetics and genomics. Sci. Adv. 9, eade1204 (2023).

36. Mollandin, F. et al. Guiding eQTL mapping and genomic prediction of gene expression in three pig breeds with tissue-specific epigenetic annotations from early development. Genomics 118, 111158 (2025).

37. Sosnicki, A., Gonzalez, J., Fields, B. & Knap, P. A review of porcine skeletal muscle plasticity and implications for genetic improvement of carcass and meat quality. Meat Sci. 219, 109676 (2025).

38. Yin, H. et al. Multi-dimensional annotation of porcine variants using genomic and epigenomic features in pigs. BMC Biol. 23, 188 (2025).

